# HADEG: A Curated Hydrocarbon Aerobic Degradation Enzymes and Genes Database

**DOI:** 10.1101/2022.08.30.505856

**Authors:** Jorge Rojas-Vargas, Hugo G. Castelán-Sánchez, Liliana Pardo-López

## Abstract

Databases of genes and enzymes involved in hydrocarbon degradation have been previously reported. However, these databases specialize on only a specific group of hydrocarbons and/or are constructed partly based on enzyme sequences with putative functions indicated by *in silico* research, with no experimental evidence. Here, we present a curated database of Hydrocarbon Aerobic Degradation Enzymes and Genes (HADEG) containing proteins and genes involved in alkane, alkene, aromatic, and plastic aerobic degradation and biosurfactant production based solely on experimental evidence, which are present in bacteria, and fungi. HADEG includes 259 proteins for petroleum hydrocarbon degradation, 160 for plastic degradation, and 32 for biosurfactant production. This database will help identify and predict hydrocarbon degradation genes/pathways and biosurfactant production in genomes.

**Graphical abstract:** 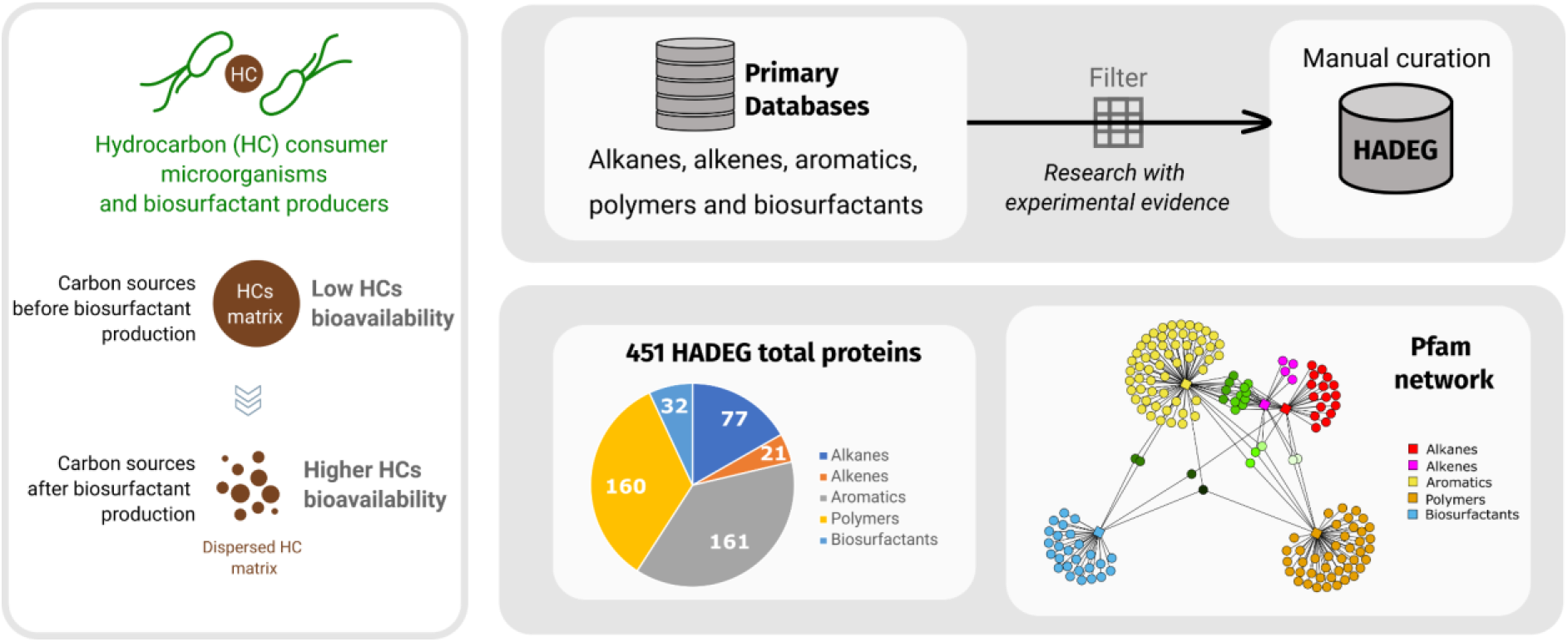

**Data summary:** The HADEG database repository is https://github.com/jarojasva/HADEG. All Supplementary Material file is available on: https://figshare.com/articles/dataset/Supplementary_Material_HADEG/20752642.

## 1 Introduction

Crude oil spills and plastic pollution are two major concerns in the context of environmental equilibrium and conservation (Bhattacharjee & Dutta, 2022; Horton, 2022). Bioremediation technologies based on microorganisms that can use hydrocarbons (HCs) and plastics as carbon sources remain among the most promising options for crude oil and plastic cleanup. Interest in bioremediation has led to the identification and characterization of some proteins involved in degradation processes. Among the features of these proteins, studies have described their activity conditions, substrates, and products, in addition to their protein structures and active sites in some cases (Brzeszcz & Kaszycki, 2018; Kaushal et al., 2021; Pathak et al., 2022; Xu et al., 2018). Biosurfactant production proteins have also been studied and shown to be involved in HCs degradation; these microbial surface-active compounds increase the bioavailability of HCs and therefore promote higher levels of HC degradation (Banat, 1995; Kavitha & Bhuvaneswari, 2021; Vimala & Mathew, 2016).

Advances in sequencing technologies have allowed the nucleotide sequences of hydrocarbon-degrading genes and, thus, the amino acid sequences of the encoded proteins to be obtained. Some of these sequences can be found in existing tailored data repositories and can be used to identify and predict the degradation potential of microbial genomes and metagenomes. Regarding hydrocarbon degradation, the AromaDeg database is focused on the aerobic degradation of aromatic compounds. It was constructed based on a phylogenomic approach and considers different oxygenases with known and unknown aromatic substrates (Duarte et al., 2014). A more recent database is that of the Calgary approach to ANnoTating HYDrocarbon degradation genes (CANT-HYD), which contains genes involved in aerobic, anaerobic aliphatic and aromatic hydrocarbon degradation pathways. CANT-HYD uses Hidden Markov Models (HMMs) for the annotation of six monophyletic clades of genes related to aliphatic aerobic degradation (*alkB, almA, ladA, bmo*, CYP153, *prm*), six related to aromatics (*dmp, dsz*, MAH, *ndo, tmo, tom*), and eight related to anaerobic degradation pathways (Khot et al., 2021). The workflow derives 37 HMMs from experimental and *in silico* derived enzymes and is designed for marker gene identification and annotation, excluding some hydrocarbon degradation pathways. Neither AromaDeg nor CANT-HYD databases contain sequences related to alkene degradation.

For plastics, the Plastics Microbial Biodegradation Database (PMBD) is a resource containing 79 experimentally predicted and ∼8000 *in silico*-predicted sequences of proteins, from the literature and the UniProt database (Gan & Zhang, 2019). The Plastics-Active Enzymes Database (PAZy) is an inventory of enzymes that act on only four synthetic fossil fuel-based polymers and three polymers mainly found in renewable resources (Buchholz et al., 2022). Additionally, PlasticDB (Gambarini et al., 2022) includes 174 experimentally tested proteins (consulted on July 15, 2022) related to the degradation of seven natural and twenty synthetic plasctics. Regarding biosurfactants, the only reported database for these compounds is BioSurfDB (www.biosurfdb.org) (Oliveira et al., 2015), which includes some experimental and predicted proteins.

Here, we introduce the manually curated database of Hydrocarbon Aerobic Degradation Enzymes and Genes (HADEG), which is a public repository containing sequences of experimentally characterized proteins and genes from bacteria and fungi. This comprehensive database is a valuable resource for the annotation of genes with potential HC degradation functions. HADEG groups these molecules according to their substrates (in the case of alkane, alkene, aromatic, and plastic aerobic degradation) and products (in the case of biosurfactant production). In this work, we also briefly reviewed the proteins and pathways involved in petroleum hydrocarbon and plastic degradation.

## 2 Materials and Methods

### 2.1 Selection of reference sequences

For proteins involved in alkane, alkene, and aromatic aerobic degradation, we compiled and classified metabolic pathways from the primary literature and two experimentally validated curated databases, MetaCyc (Krieger et al., 2004) and KEGG (Ogata et al., 1999). We retrieved the sequences of proteins and genes involved in the aforementioned metabolic pathways in the well-recognized repository for experimentally characterized proteins UniProtKB/Swiss-Prot database (Boutet et al., 2007), and in the literature (Supplementary Tables 1, 2, and 3). We used the NCBI portal (https://www.ncbi.nlm.nih.gov/protein) to obtain the sequences with their accession numbers if they were not available in any database.

To obtain plastic degradation proteins, we selected nonredundant protein sequences from PlasticDB (Gambarini et al., 2022). To obtain biosurfactant production proteins, we used the UniProt platform with different biosurfactants as keywords, and the corresponding proteins included in the UniProtKB/Swiss-Prot database were collected. In constructing our plastic and biosurfactant database, we added some new proteins from a search of the primary literature (Supplementary Tables 4 and 5). The nucleotide sequences of all these proteins were retrieved from the European Nucleotide Archive (ENA) platform (Leinonen et al., 2011) or the literature.

### 2.2 Protein annotation and orthology

We used eggNOG-mapper (Huerta-Cepas et al., 2019) online version 2.1.9, run on July 23 (2022), with default parameters, to annotate our protein database, identify orthologs, and study their taxonomic distribution at the phylum level. Protein sequences without associated gene orthologs were considered as seed sequences. We used InterProScan v.5.51-85.0 (Jones et al., 2014) for protein domain inference with the default parameters to identify the shared protein domains predicted with Pfam-33.1 (Supplementary Table 7).

### 2.3 Searching for hydrocarbon degradation proteins in genomes using HADEG

We selected 66 genomes of recognized HC and plastic-degrading bacteria and fungi, and biosurfactant producers for validation (Supplementary Table 8). In addition, four genomes from anaerobic bacteria and one genome from a non-degrading fungus were selected as negative controls, for a total of 71 genomes (Supplementary Table 8). The FASTA files of the proteins were retrieved from the NCBI portal (accessed June 23, 2023) and annotated with KofamKOALA v2023-04-01 (Aramaki et al., 2020) to obtain the KO codes of each protein. Proteinortho v6.1.7 (Lechner et al., 2011) was used to predict orthologous proteins by applying different identity cutoffs (25, 50, 75) and minimal algebraic connectivity (0.0, 0.1, 0.2, 0.3, 0.4) against the protein FASTA files from our database. After testing these ranges of values, functional annotations were considered correct if the KO codes matched between HADEG proteins and the identified orthologs in the genomes. False-negative hits were determined by searching for 125 expected hits among the genomes (Supplementary Table 9) using the proteins with correct annotations. The “identity” and “conn” values that minimized the number of false-negative hits and provided at least 90% of correct annotations were selected.

### 2.4 Comparison of HADEG with other databases

The HADEG pathways were compared to the pathways included in the specialized databases CANT-HYD, AromaDeg, PAZy, PlasticDB, and BioSurfDB. The presence or absence of pathways was examined using a heatmap and an upset plot with the library ComplexHeatmap in R (Gu et al., 2016).

## 3 Results and Discussion

### 3.1 Hydrocarbon aerobic degradation

HCs are relatively stable molecules but can serve as a source of carbon and energy for any organism able to activate them (Widdel & Musat, 2010). This activation by the aerobic insertion of oxygen or the anaerobic addition of succinate distinguishes hydrocarbon-degrading organisms from their counterparts (Prince et al., 2010). In HADEG, we included proteins related to aerobic HC degradation, such as alkanes, alkenes, aromatics, and plastics (Supplementary Tables 1-4).

#### 3.1.1 Alkanes

Alkanes are the principal components of crude oil. Four microbial pathways for alkane degradation have been identified (Fig. 1) (Ji et al., 2013; Moreno & Rojo, 2017; Wentzel et al., 2007). All of them begin with the activation of the alkane chain through its oxidation to a primary or secondary alcohol, as observed in terminal (TO), biterminal (BO), and subterminal oxidation (SO), or to an *n*-alkyl hydroperoxide via the Finnerty pathway (FP). Among the activation proteins included in HADEG, we report the extensively studied alkane monooxygenase (*alkB*), cytochrome P153 (CYP153), and P450 (CYP450), flavin-binding protein monooxygenase (*almA*), long-chain alkane monooxygenase (*ladA*), and terminal alkane-hydroxylase (*alkM*). The rubredoxins (*rubA, rubB*), which are crucial in electron transfer to *alkB*, are also included. Some other included proteins belong to particular pathways, such as Baeyer-Villiger monooxygenase (BVMO) in SO and alkyl hydroperoxide reductase (*ahpCF*) in FP. Proteins with specific substrates, such as particulate methane monooxygenase (*pmoA*), propane 2-monooxygenase (*prm*), and butane monooxygenase (*bmo*), are also included in HADEG.

**Fig. 1.**
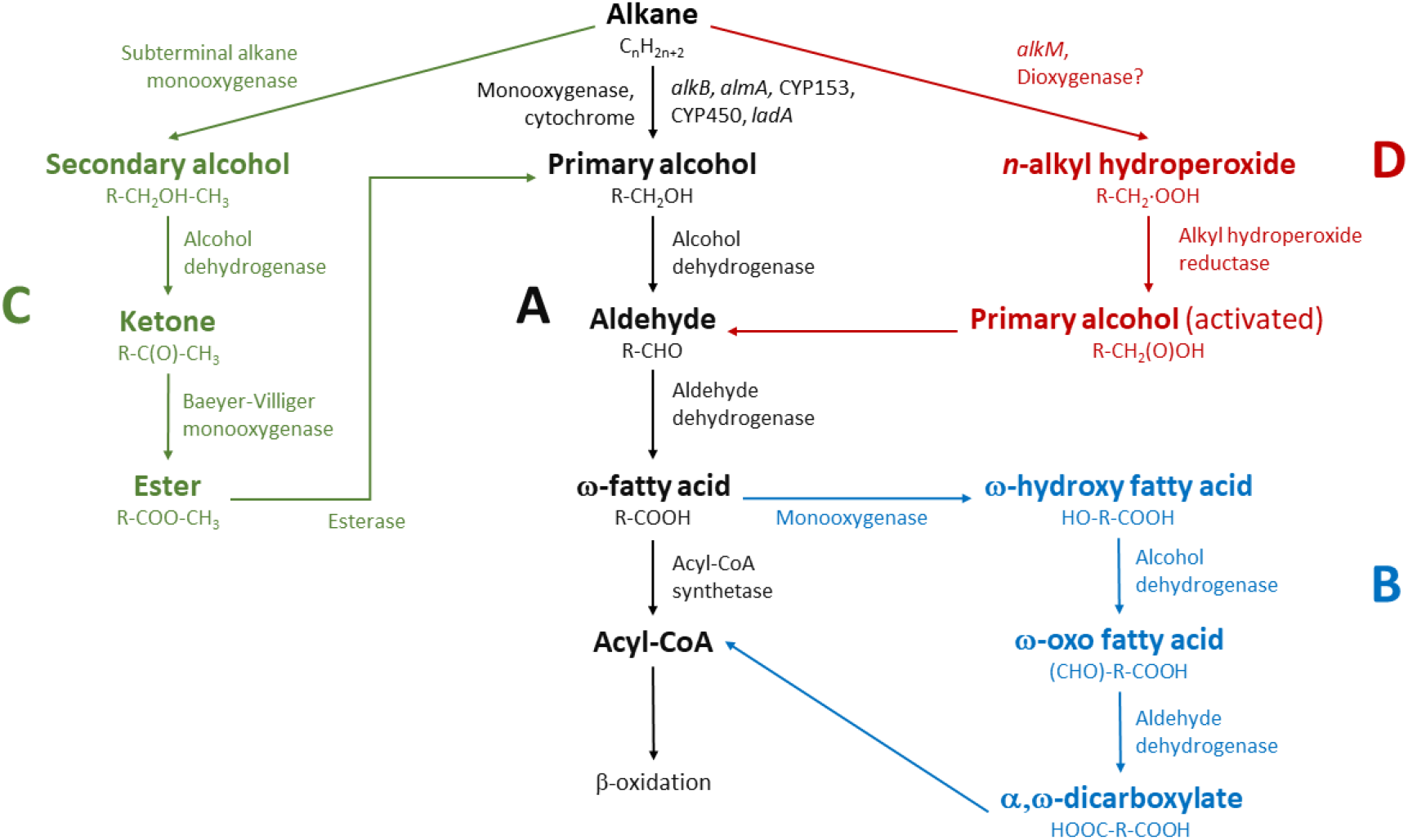
Alkane aerobic degradation pathways. **A**. Terminal oxidation (black color). **B**. Biterminal oxidation (blue). **C**. Subterminal oxidation (green). **D**. Finnerty pathway (red).

Other proteins included in HADEG are alcohol and aldehyde dehydrogenases (*adh, ald*), cyclohexanone 1,2-monooxygenase (CHMO), alkanesulfonate monooxygenase (*ssuD*), the helix-turn-helix transcriptional regulator of *alkB* (*alkS*), and enzymes related to the uptake of alkanes, such as outer membrane protein (*alkL*), outer membrane lipoprotein (*blc*), long-chain fatty acid transport enzyme (*fadL*), and methyl-accepting chemotaxis protein (*alkN* or *mcp*). All the proteins for alkane degradation included in HADEG are described in Supplementary Table 1.

#### 3.1.2 Alkenes

Alkenes are transformed into the corresponding epoxides by alkene monooxygenases. Epoxides are somewhat toxic to cells but can be processed using epoxide hydrolases, such as the protein epoxyalkane CoM transferase, and their products ultimately enter the central metabolism. HADEG includes the alkene monooxygenase complex (*amoABCD*) identified in *Nocardia corallina* B-276, the isoprene monooxygenase (*isoABCDEHI*) from *Rhodococcus* sp. AD45, the propene monooxygenase system (*xamoABCDE*) and the epoxyalkane CoM system (*xecADC*) from *Xanthobacter autotrophicus* Py2, the epoxyalkane CoM (*etnE*) from *Mycolicibacterium rhodesiae*, and the dehydrogenase *mpdBC* from *M. austroafricanum*. All the proteins for alkene degradation included in HADEG are described in Supplementary Table 2.

#### 3.1.3 Aromatics

Aromatic compounds are commonly known as recalcitrant pollutants because of their relatively low susceptibility to microbial degradation (Ghosal et al., 2016). Some also present mutagenic and carcinogenic risks to human health. In contrast to alkanes and alkenes, aromatic compound degradation requires complex metabolic machinery. In general, aerobic organisms take advantage of the availability of molecular oxygen and use oxygenases that introduce hydroxyl groups and cleave aromatic rings. These pathways usually involved a few central intermediates, most commonly including catechol, protocatechuate, and gentisate (Phale et al., 2020).

The goal of degradation is the activation and cleavage of aromatic rings to obtain products that can subsequently be integrated into central pathways. Our research included pathways with characterized proteins involved in the metabolic processes of poly- and monoaromatic degradation. Regarding polyaromatic degradation pathways, HADEG includes proteins involved in the degradation of anthracene, biphenyl, dibenzothiophene, fluorene, naphthalene, and phenanthrene. Regarding monoaromatic degradation pathways, we included proteins involved in the degradation of benzene, toluene, xylene (BTX), anthranilate, benzoate, 2-nitrobenzoate, catechol (ortho- and meta-cleavage pathways), gentisate, phenol, phenylacetate, phthalate, protocatechuate (ortho- and meta-cleavage pathways), salicylate, styrene, and terephthalate. All the proteins for aromatic degradation included in HADEG are described in Supplementary Table 3.

#### 3.1.4 Plastics

Proteins involved in the degradation of plastics were also added to the HADEG database, since plastics are important environmental contaminants, and it has been reported that some microorganisms can degrade them (Kaushal et al., 2021). Some plastics are commonly derived from petroleum (e.g., polyethylene terephthalate-PET, polyurethane-PU, polyvinyl chloride-PVC, polystyrene-PS, and polypropylene-PP) and have many applications (Shrivastava, 2018). Most plastics are very difficult to degrade due to inherent characteristics such as high molecular weights, crystallinity, and hydrophobicity (Amobonye et al., 2021; Geyer, 2020). In this context, microorganisms capable of plastic degradation remain one of the most promising options for reducing or recycling plastics with long lifetimes (Muriel-Millán et al., 2021).

The plastic degradation process begins with superficial deterioration and subsequent cleavage of plastics into intermediates, which can be used as energy and carbon sources. The depolymerization process mainly involves hydrolytic cleavage of glycosidic, ester, and peptide bonds, resulting in constituent oligomers or monomers (Amobonye et al., 2021). Implementing physicochemical pretreatments such as the application of chemical additives or UV irradiation can facilitate microbial plastic degradation (Ru et al., 2020).

Among the plastic degradation-proteins included in HADEG, there are some hydrolases for PET degradation, such as the recently well-characterized polyester hydrolase of *Halopseudomonas aestusnigri* VXGO4 (6SCD PDB), the GEN0105 esterase from a marine metagenome (Hajighasemi et al., 2018), a lacasse of *Rhodococcus ruber* C208, and some cutinases. These enzymes also include PVA dehydrogenase (*pvaA, pvaB*) for polyvinyl alcohol (PVA) degradation (Shimao, 2001); exo-cleaving and endo-cleaving rubber dioxygenases (*roxA, roxB*) for natural rubber degradation from *Xanthomonas* sp. 35Y; polyurethanases (*pueA, pueB*) for PU degradation from *Pseudomonas chlororaphis* and *Pseudomonas protegens* Pf-5; manganese peroxidase 1 (*mnp1*) for polyethylene degradation from *Phanerodontia chrysosporium*; and a cutinase for poly(epsilon-caprolactone) (PCL) and poly(1,3-propylene adipate) (PPA) degradation from *Kineococcus radiotolerans* DSM 14245. All the proteins for plastic degradation included in HADEG are described in Supplementary Table 4.

### 3.2 Biosurfactants production

Including sequences for biosurfactant production in HADEG will greatly enhance the database’s ability to provide a comprehensive overview of the HC-degrading potential of microorganisms. Biosurfactants are surface-active compounds that play a crucial role in the initial steps of HC degradation by facilitating access to HC molecules. Some microorganisms capable of degrading HCs secrete biosurfactants of different natures, such as lipoproteins or rhamnolipids, to emulsify HCs.

Previous reports and reviews (Baldi et al., 1999; Hua & Wang, 2014; Moreno & Rojo, 2017) have discussed the observation that short-chain alkanes containing fewer than 9 carbons are sufficiently soluble to be easily transported into microbial cells. However, for other HC compounds, microorganisms need to directly adhere to oil droplets in order to access the target hydrocarbons. Biosurfactants are believed to play a crucial role in this process by increasing the surface area available for hydrophobic compounds to interact with the aqueous phase. This increase in surface area facilitates the access of microorganisms to the oil phase. Additionally, studies have observed that biosurfactants can increase the surface hydrophobicity of cells and enhance membrane permeability (Kaczorek et al., 2018). These effects further contribute to the improved interaction between microorganisms and the hydrophobic compounds, ultimately facilitating their utilization by microorganisms.

The HADEG database includes sequences of genes and proteins involved in the production of fengycin (*ffp*), plipastatin (*ppsABCDE*), surfactin (*srf*), iturin (*itu*), and mycosubtilin (*mycABC*) in *Bacillus subtilis*. Additionally, genes/proteins related to rhamnolipids (*rhl*) from *Pseudomonas aeruginosa*; emulsan (*wzb, wzc*) from *Acinetobacter lwoffii* RAG-1; arthrofactin (*arfABC*) from *Pseudomonas* sp. MIS38; amphisin (*amsY*) from *Pseudomonas* sp. DSS73; serrawetin (*pswP*) in *Serratia marcescens*; and lactonic sophorose lipid (*sble*), and sophorolipids (*at*) from *Starmerella bombicola* are also included. Two fungal biosurfactants are also included; mannosylerythritol lipids (*emt1*) from *Ustilago maydis* and hydrophobin (*hfb1, hfb2*) from *Hypocrea jecorina*. All the proteins for biosurfactant production included in HADEG are described in Supplementary Table 5.

### 3.3 Orthologs of the HADEG database

Among the protein sequences present in HADEG, orthologs were searched to determine their distribution in different microorganisms and their taxonomic assignments. We used eggNOG-mapper to identify 19,749 orthologs of 445 protein sequences among the 451 sequences included in HADEG (Supplementary Table 6). The six proteins without identified orthologs were the PBAT esterase of an uncultured bacterium (accession no. A0A1C9T7G6), the PVA hydrolase of *Sphingomonas sp*. (accession no. Q588Z2, *oph*), the butane monooxygenase hydroxylase gamma subunit of *Thauera butanivorans* (a.n. Q8KQE7, *bmoZ*), the fluoren-9-ol dehydrogenase of *Terrabacter* sp. DBF63 (a.n. Q93UV4, *flnB*), the oxidized PVA hydrolase of *Pseudomonas* sp. (a.n. Q9LCQ7, *pvaB*), and the anthranilate-CoA ligase of *Rhodococcus erythropolis* (a.n. S6BVH5, *rauF*).

The taxonomic analysis of orthologs at the phylum level revealed that Proteobacteria was the most abundant phylum related to HC degradation pathways and biosurfactant production (forest-green color in Fig. 2A). Many studies have confirmed the degradation capacities of species in genera such as *Alcanivorax* and *Marinobacter* (hydrocarbonoclastics), *Acinetobacter* (versatile HC degraders), and ubiquitous *Pseudomonas*. Biosurfactants such as amphisin, arthrobactin, emulsan and serrawetin have only been described in Proteobacteria (Soberón-Chávez, 2010), and rhamnolipid production mechanisms have been well described in *Pseudomonas* (Soberón-Chávez et al., 2021) (Fig. 2B). Actinobacteria was the second most abundant phylum related to HC degradation pathways (goldenrod color in Fig. 2), and it includes the hydrocarbon-degrading genera *Rhodococcus, Mycobacterium, Norcadia, Gordonia*, and *Streptomyces*, among others. Firmicutes was the third most abundant phylum related to HC pathways and the second most abundant related to biosurfactant production (*Bacillus, Geobacillus*, and *Phenyracillin*) (royal blue color in Fig. 2), within which surfactin and iturin production by *Bacillus* genus members has been well studied (Soberón-Chávez, 2010). Khot et al. (2021) also identified these three phyla with the higher number of hits across the CANT-HYD HMMs for alkane and aromatic degradation. Among the Bacteria domain, the fourth most abundant phylum was Bacteroidetes (*Aequorivita, Algoriphagus, Cytophaga*), particularly related to alkene degradation (orange color in Fig. 2). Ascomycota was the most abundant phylum from the fungal domain and was mainly associated with alkane, and plastic degradation and biosurfactant production (brick color in Fig. 2) (see Supplementary Table 6). Fungi have received interest in relation to bioremediation applications because they do not rely solely on soluble organic compounds for nutrition; they also show diverse enzymatic mechanisms and distinct mechanisms of physiological adaptability to environmental conditions (Prenafeta-Boldú et al., 2018).

**Fig. 2.**
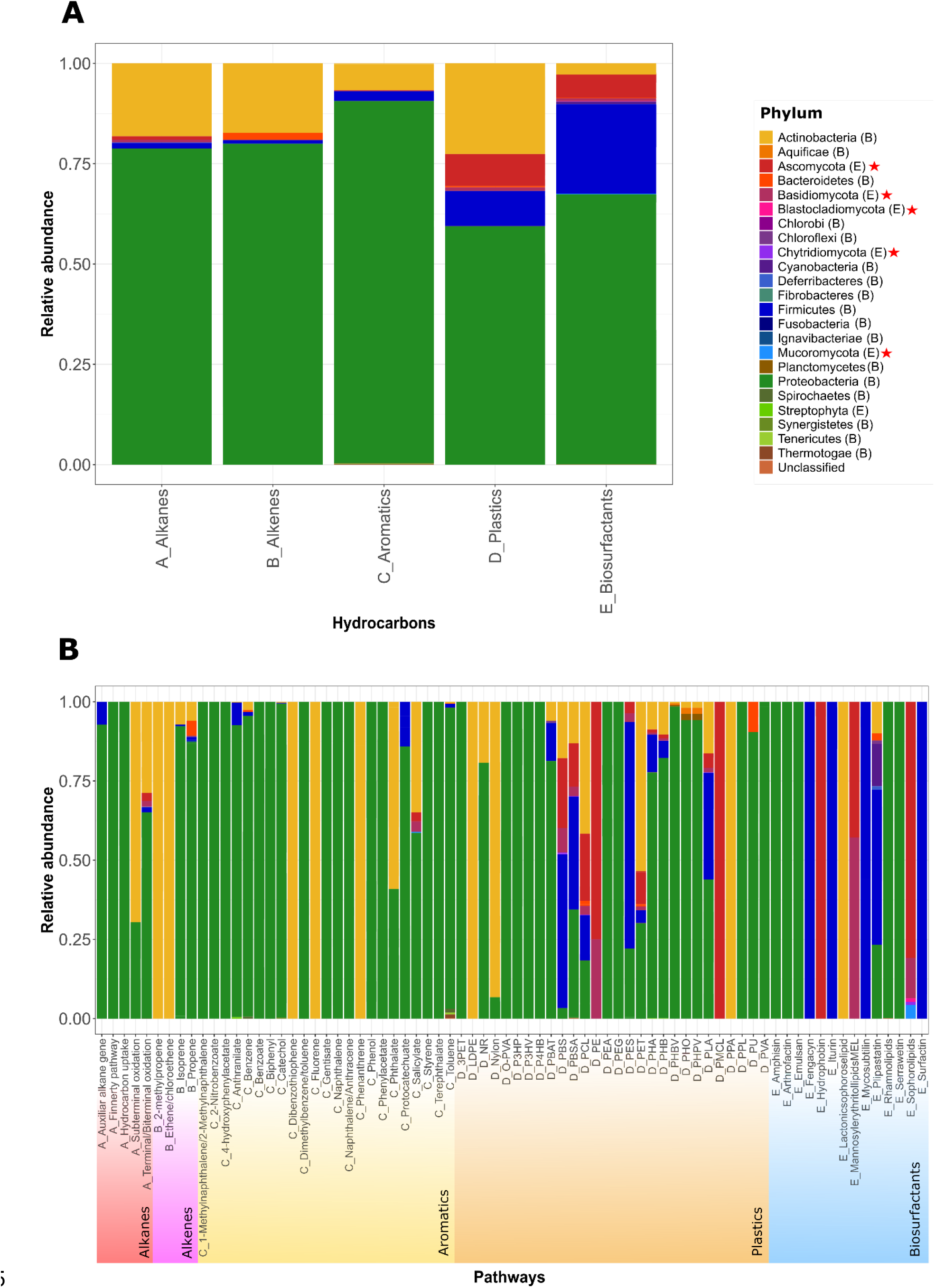
Taxonomic distribution of orthologous proteins determined with eggNOG mapper. **A**. hydrocarbon groups. **B**. Degradation pathways. A_alkane groups, B_alkenes, C_aromatics, D_plastics, E_biosurfactants. Fungal phyla have a red asterisk in the legend. (B) represents bacterial phyla, (E) represents eukaryotic phyla.

### 3.4 HADEG Pfams network

A network analysis was performed to determine whether the HADEG proteins shared domains, which may indicate that a protein could participate in the degradation of more than one type of HC (Fig. 3). Identifying the shared and nonshared protein domains between hydrocarbon degradation groups will be helpful to establish a set of biomarkers for the targeted searching of potential hydrocarbon-degrading microorganisms. A total of 166 Pfam domains were identified in HADEG (Supplementary Table 7). Network analyses reveal that the plastic and aromatic degradation groups contained the largest numbers of protein domains, at 75 and 48, respectively. These results are congruent with the diversity and complexity of aromatic and plastic chemical structures. Additionally, we predicted 30 Pfams among biosurfactants, 28 among alkanes, and 18 among alkenes.

**Fig. 3.**
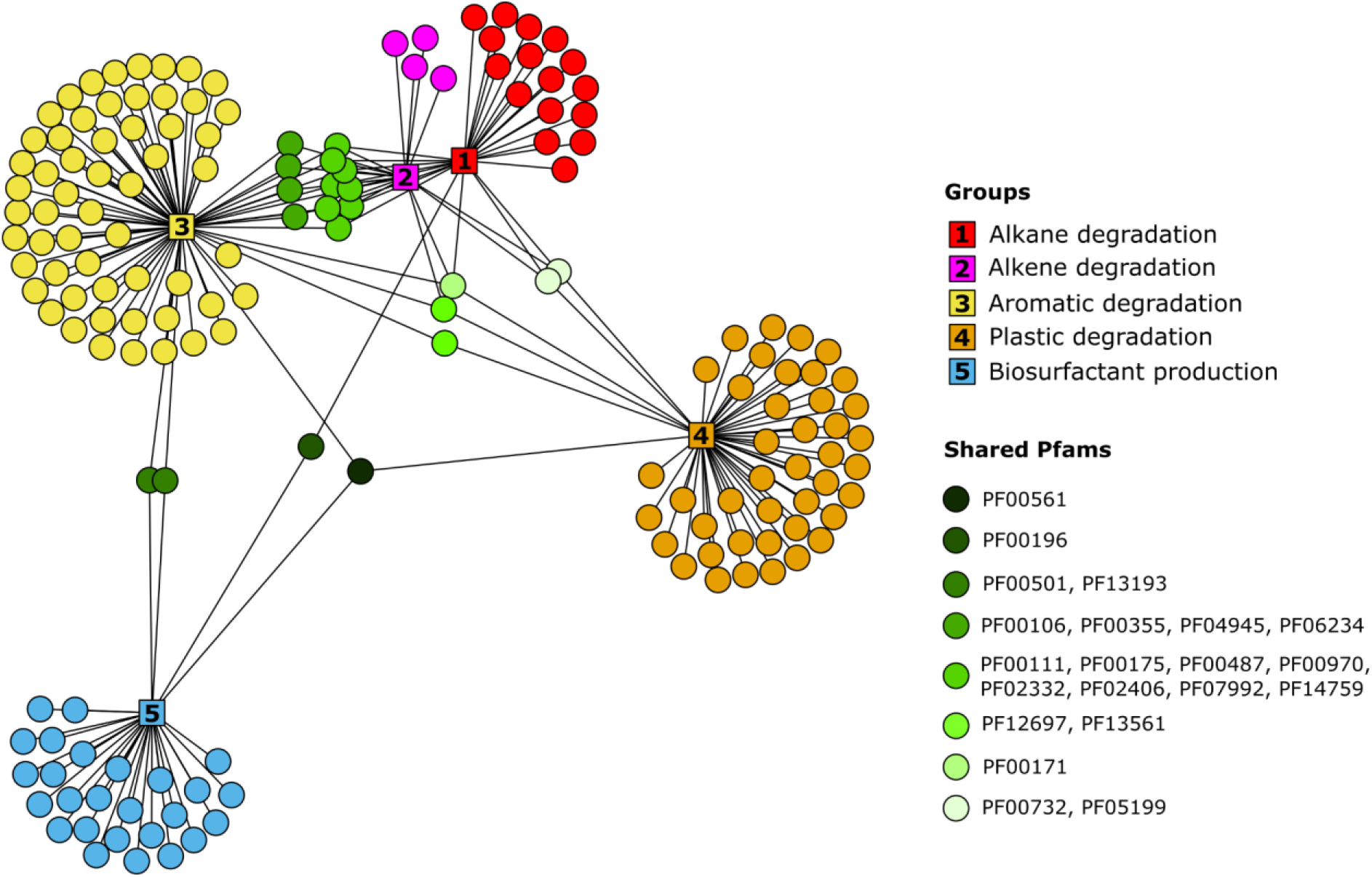
HADEG Pfams network. The circles represent the Pfams: for alkane degradation in red, for alkene degradation in magenta, for aromatics degradation in yellow, for plastic degradation in orange, and for biosurfactant production in blue. The shared protein domains among groups are shown on a green scale. For a complete list of Pfams, see Supplementary Table 7.

Among the protein domains without connectivity or non-shared protein domains, the alkane set included 16 Pfams such as the related to NAD(P)-binding Rossmann-like domain (PF13450) and a flavin-binding monooxygenase-like protein (PF00743) in *almA* and BVMO. The alkene set included four Pfams with no connectivity: cobalamin-independent synthase (PF01717) in *etnE*, a glutathione S-transferase C-terminal domain (PF17171 and PF17172) in *isoI*, and a pyridine nucleotide-disulfide oxidoreductase dimerization domain (PF02852) in *xecC*. The aromatic set included 57 nonshared Pfams like the ring-hydroxylating alpha and beta subunits (PF00848 and PF00866) present in aromatic-ring dioxygenase systems, the dioxygenase superfamily (PF00903), and the catechol dioxygenase N-terminus (PF04444). The plastic set included 42 nonshared protein domains like the RTX calcium-binding nonapeptide repeat 4 copies (PF00353) present in the polyester polyurethanases *pueA* and *pueB*, the alpha/beta hydrolase family (PF12695) involved in PET degradation, the esterase PHB depolymerase (PF10503) involved in PHA/PHB degradation and a cutinase (PF01083) that acts on carboxylic ester bonds. Finally, the biosurfactant set included 26 nonshared Pfams like a condensation domain (PF00668) present in *Bacillus* biosurfactants synthesis proteins (iturin, mycosubtilin, surfactin, and plipastatin), the 4’-phosphopantetheinyl transferase superfamily (PF01648) in *ffp, pswP* and *sfp*, and helicase domains (PF00270 and PF00271) in *rhlB* and (PF13641) in *rhlC*, both involved in rhamnolipids production.

Concerning the protein domains that were connected in the network or shared, only the aldehyde dehydrogenase family (PF00171) participates in the degradation of the four HC groups. This domain is responsible for the oxidation of aldehydes, which is considered a detoxification reaction (*Pfam: Family: Aldedh (PF00171)*, n.d.). The largest set of shared domains was found between alkenes and aromatics, which shared 19 Pfams. These domains included the Rieske [2Fe-2S] domain (PF00355), which is involved in the electron transfer chain. PF00355 was present in enzymes such as *andA, benA, doxAB, ndoAB, tmoC, todB, bedB, isoC*, and *xamoC*. The alkane, alkene, and aromatic groups shared seven Pfam domains, including the oxidoreductase NAD-binding and FAD-binding domains (PF00175 and PF00970), present in *prmB, isoF, xamoF, benC*, and *tmoF*; the methane/phenol/toluene hydroxylase domains (PF02332), found in *bmoX, xamoA*, and *tmoA*; and the MmoB/DmpM family (PF02406), found in *prmD, xamoD*, and *tmoD*. The alkane and aromatic groups shared a fatty acid desaturase domain (PF00487) that catalyzes the insertion of a double bond at the delta position in fatty acids (*Pfam: Family: FA_desaturase (PF00487)*, n.d.). This domain is present in alkane monooxygenases (*alkB*), alkane hydroxylases (*alkMa, alkMb*), and xylene monooxygenase subunit 1 (*xylM*).

### 3.5 Hydrocarbon degradation proteins in different genomes

The HADEG database is a valuable repository for the identification of HC degradation proteins in genomes. In this study, we used an orthology-based identification strategy with the Proteinortho program. Proteinortho uses a graph representation of proteins and their similarities determined by BLAST to identify orthologous proteins between genomes. Spectral partitioning techniques are then applied to identify groups of orthologous proteins or proteins with similar functions, allowing for the identification of orthologs across genomes of closely related or distantly related species (Lechner et al., 2011).

A Proteinortho analysis was performed comparing HADEG with protein FASTA files from 71 genomes (refer to Supplementary Table 8 for the complete list of genomes). Orthologous groups were inferred by adjusting different minimum sequence identities and algebraic connectivities for spectral partitioning. We then assessed the number of Predicted Orthologous Proteins (POPs) with similar KO annotations to those from HADEG. Our results revealed that using the parameters -identity=50 and -conn=0.3 resulted in the fewest false-negative hits (26 out of 125 expected hits, 20.80%) and a high correct annotation of 94.68% of POPs (1,335 out of 1,410) (see Supplementary Table 10). An identity of 50% and a connectivity of 0.1, produced an equal number of false-negative hits, but the percentage of correct annotations was lower (92.42%). Increasing the minimum identity threshold to 75% resulted in more than 99% of POPs being correctly annotated, but also resulted in a higher number of false-negative hits (∼64), indicating the loss of ∼51% of expected hits. Based on these results, we determined that the -identity=50 and -conn=0.3 parameters were most appropriate for identifying POPs between HADEG and the selected genomes.

Using these parameters, the analysis revealed that 65 of the 66 genomes used for validation purposes exhibited POPs that corresponded to experimentally verified metabolisms (Fig. 4). Among the genomes analyzed, the six selected fungal genomes of *Exophiala, Yarrowia, Aspergillus*, and *Fusarium* had POPs associated with genes involved in the degradation of alkanes and aromatics, such as BVMO, CHMO, and CYP450, as well as esterases, cutinases, and PHB depolymerases for plastic degradation. These fungal genera have been reported as degraders of alkanes, monoaromatics, and/or plastics (Ekanayaka et al., 2022; Madzak, 2021; Srikanth et al., 2022; Yoshida et al., 2010).

**Fig. 4.**
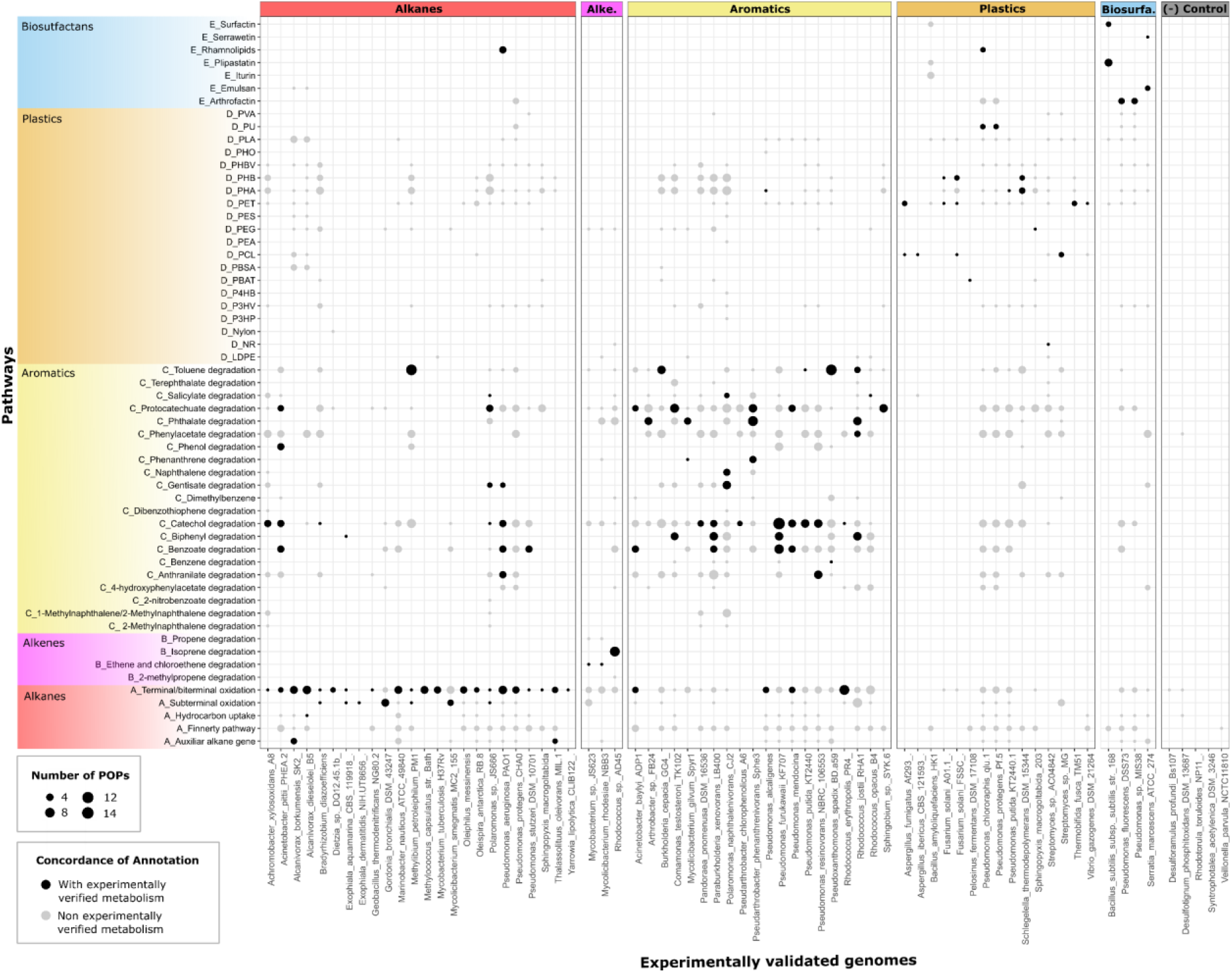
Annotation of genomes using Proteinortho against the HADEG database. Each bubble represents the number of Predicted Orthologous Proteins (POPs) with similar KO annotations to those from HADEG. The genomes (x-axis) are plotted against the hydrocarbon degradation pathways or biosurfactant production of HADEG (y-axis). The hits with experimental evidence are in black, and the potential ones are in gray. The size of the bubble represents the number of hits in the genome. A complete list of genomes and their accession number can be found in Supplementary Table 8.

The highest number of POPs (14) was observed in the catechol degradation pathway of the *Pseudomonas furukawaii* KF707 assembly. These POPs were associated with genes for the ortho-cleavage mechanism (*catA, catB, catC, pcaD, pcaF*), as well as for the meta-cleavage mechanism (*xylE, xylH, xylF*, and two copies of *xylG, xylI, xylJ*). Strain KF707 uses both cleavage mechanisms (Murphy et al., 2008). The second highest number of POPs (11) was found in the toluene degradation pathway of the *Methylibium petroleiphilum* PM1 genome. These POPs were linked to genes of the toluene-4-monooxygenase complex (*tmoA, tmoB, tmoC, tmoD, tmoE*) and the toluene ortho-monooxygenase complex (*tom1, tom3, tom4, tom5*, and two copies of *tom2*). However, the specific complex involved in toluene degradation in PM1 remains unknown (Deeb et al., 2001). The only genome lacking expected POPs belonged to the strain *Bacillus amyloliquefaciens* HK1, despite experimental evidence of its capability to degrade PVA (Zhang et al., 2018). This result suggested that this isolate likely has proteins involved in PVA degradation that are not currently present in our database.

Our analysis identifies unreported HC degradation and biosurfactant production capabilities in 60 of the 66 genomes. Many of these genomes included POPs associated with genes for aliphatic, aromatic, and plastics degradation and the production of biosurfactants. The occurrence of these numerous pathways might be due to microbial environmental adaptations and be widely observed within the microbial world (Brzeszcz & Kaszycki, 2018). However, empirical evidence suggests that microbes show metabolic pathway preferences (Brzeszcz & Kaszycki, 2018).

Using Proteinortho, we also detected five POPs in three negative control genomes, a very reduced number compared with the predicted for the other genomes. These POPs were associated with rubredoxin reductase (*rubB*), outer membrane lipoprotein (*blc*), phenylacetate-coenzyme A ligase (*paaK*), and alkyl hydroperoxide reductase subunit C (*ahpC*). These proteins are known to be involved in basic cellular functions such as electron transfer chain, cell homeostasis regulation, and detoxification processes. The annotation of these POPs was consistent with the NCBI Prokaryotic Genome Annotation Pipeline.

The concordance between experimental evidence and the predicted pathway repertoire demonstrates that the HADEG database is a reliable sequence database to be used to predict the potential HC degradation pathways and biosurfactant production in genomes.

### 3.6 Comparison of HADEG with other databases

We compared HADEG with several databases including CANT-HYD, AromaDeg, PAZy, PlasticDB, and BioSurfDB. Our analysis involved examining the metabolic pathways within each database and visualizing them using a heatmap and an upset plot (Fig. 5).

**Fig. 5.**
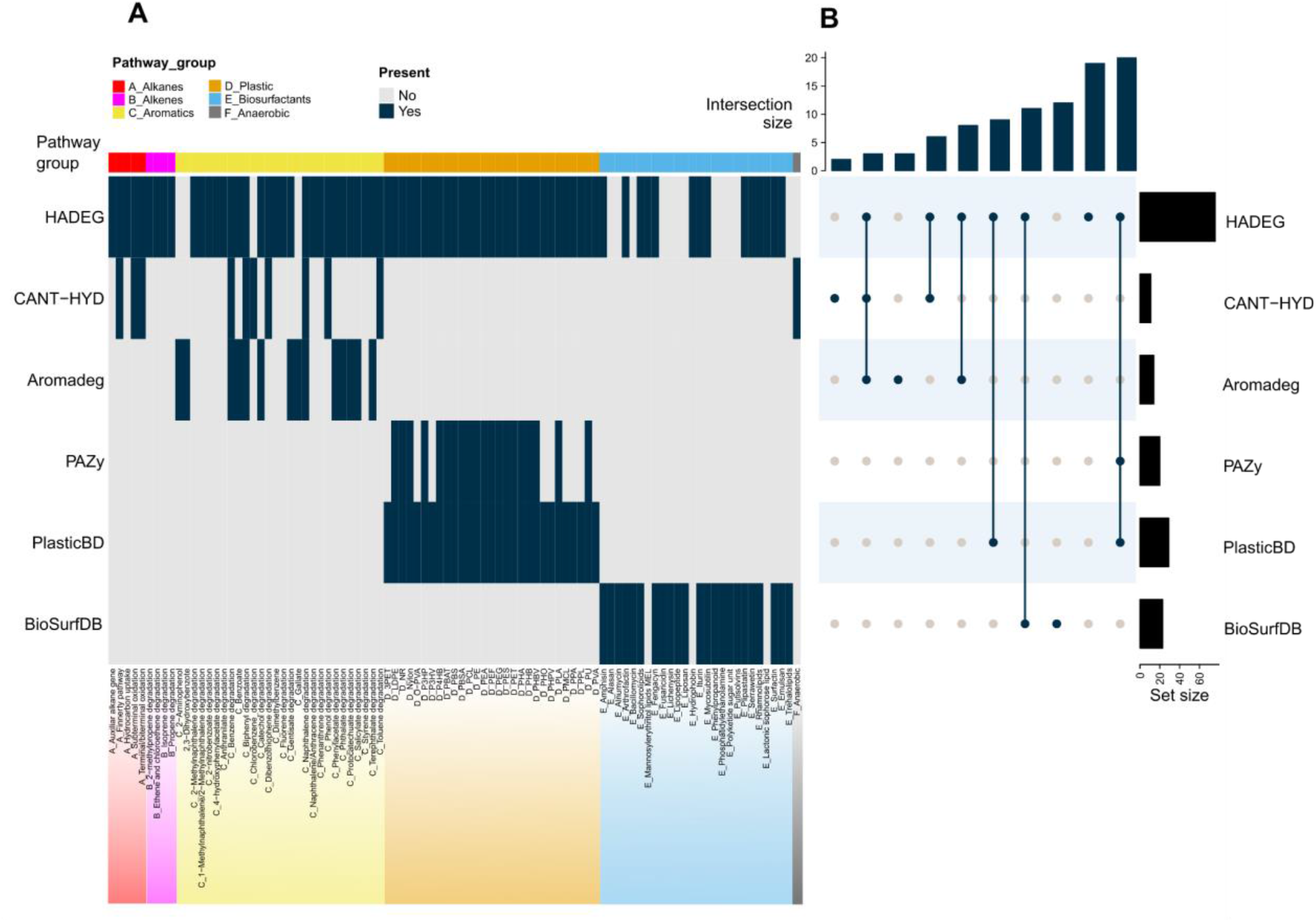
Comparison of HADEG with databases CANT-HD, PAZy, PlasticDB, Aromadeg, and BioSurfDB. **A**. Heatmap of metabolic pathways present in each database. HADEG showed a larger number of metabolic pathways involved in HC degradation for which there is experimental evidence. **B**. Upset plot of common and unique pathways.

The results demonstrate that HADEG is a more comprehensive database, as it covers aerobic degradation pathways for alkanes, alkenes, aromatics, plastics, and production of biosurfactants (Fig. 5A). When comparing HADEG with CANT-HYD, CANT-HYD has a narrower scope, focusing on aerobic alkane and aromatic degradation, as well as anaerobic degradation pathways. In contrast, HADEG incorporates a broader range of pathways for alkane and aromatic degradation. AromaDeg is dedicated to aromatic compound degradation and includes three pathways not covered by HADEG (gallate, 2-aminophenol, and 2,3-dihydroxybenzene). Nevertheless, HADEG encompasses a total of 13 more pathways compared to AromaDeg.

While PAZy and PlasticDB specifically target plastic degradation, our database covers both plastics and other HCs. HADEG shares degradation pathways for plastics with PlasticDB, whereas PAZy lacks specific plastic degradation pathways included in HADEG, such as 3PET, O-PVA, P3HV, PHO, PHPV, PMCL, PPA, PPL, and PVA. In terms of biosurfactant production, BioSurfDB includes twelve additional pathways compared to HADEG, albeit some rely on in silico predictions. HADEG features protein sequences for three additional biosurfactant pathways not included in BioSurfDB: mannosylerythritol lipids, hydrophobins, and lactonic sophorose lipids. Among the analyzed databases, HADEG stands out as the sole repository that includes alkene degradation pathways.

The implementation of an upset plot provided valuable insights into the overlapping and distinct metabolic pathways covered by all six databases (Fig. 5B). It confirmed that HADEG presents the most extensive collection of pathways (set size), followed by PlasticDB, BioSurfDB, PAZy, AromaDeg, and CANT-HYD. Furthermore, HADEG includes 19 unique pathways not found in the other databases (intersection size in Fig. 5B), including degradation pathways for ethene, isoprene, ethene, methylnaphthalene, dimethylbenzene, fluorene, anthranilate, and phenanthrene (Fig. 5A). These results demonstrate that, unlike the other databases analyzed, HADEG enables the prediction of a greater number of metabolic pathways in genomes.

## 4 Conclusions

The HADEG database is a bioinformatic resource that can be used to identify potential microorganisms for developing HC-bioremediation methods. To the best of our knowledge, HADEG, which contains 451 experimentally verified proteins, is the first database to integrate different types of HCs (alkane, alkenes, aromatics, and plastics) and biosurfactants in a single resource. Through ortholog analysis, we found that the HADEG proteins are distributed in bacteria and fungi and that they are most highly represented in Proteobacteria. Additionally, Pfam analysis showed that the domains of the HC degradation proteins correspond primarily to plastics and aromatics. Most domains are not shared among HC groups, indicating that their functions are specific to HC-degradation and that they can probably be used as biomarkers for the identification of microbes with degradation abilities. Finally, HADEG covers more metabolic pathways for HCs degradation than other specialized databases and is a reliable tool for annotating HC and plastic degradation, and biosurfactant production proteins.

## Author contributions

**Jorge Rojas-Vargas**: Conceptualization, Data Curation, Formal analysis, Methodology, Software, Visualization, Writing-Original draft preparation. **Hugo G. Castelán-Sánchez**: Data Curation, Formal analysis, Methodology, Software, Visualization, Writing-Original draft preparation, and Editing. **Liliana Pardo-López**: Validation, Resources, Writing-Reviewing and Editing. All authors approved the final version.

## Acknowledgments

We thank the Unidad de Secuenciación Masiva y Bioinformática del Instituto de Biotecnología-UNAM for giving us access to its computer cluster. We are grateful to Ph.D. Valerie de Anda and Ph.D. Ernesto Pérez-Rueda for the ortholog’s analysis guidance.

## References

Amobonye, A., Bhagwat, P., Singh, S., & Pillai, S. (2021). Plastic biodegradation: Frontline microbes and their enzymes. The Science of the Total Environment, 759, 143536. 10.1016/j.scitotenv.2020.143536

Aramaki, T., Blanc-Mathieu, R., Endo, H., Ohkubo, K., Kanehisa, M., Goto, S., & Ogata, H. (2020). KofamKOALA: KEGG Ortholog assignment based on profile HMM and adaptive score threshold. Bioinformatics, 36(7), 2251–2252. 10.1093/bioinformatics/btz859

Baldi, F., Ivošević, N., Minacci, A., Pepi, M., Fani, R., Svetličić, V., & Žutić, V. (1999). Adhesion of Acinetobacter venetianus to diesel fuel droplets studied with in situ electrochemical and molecular probes. Applied and Environmental Microbiology, 65(5), 2041–2048. 10.1128/aem.65.5.2041-2048.1999

Banat, I. M. (1995). Biosurfactants production and possible uses in microbial enhanced oil recovery and oil pollution remediation: A review. Bioresource Technology, 51(1), 1–12. 10.1016/0960-8524(94)00101-6

Bhattacharjee, S., & Dutta, T. (2022). Chapter 1 - An overview of oil pollution and oil-spilling incidents. In P. Das, S. Manna, & J. K. Pandey (Eds.), Advances in Oil-Water Separation (pp. 3–15). Elsevier. 10.1016/B978-0-323-89978-9.00014-8

Boutet, E., Lieberherr, D., Tognolli, M., Schneider, M., & Bairoch, A. (2007). UniProtKB/Swiss-Prot. In D. Edwards (Ed.), Plant Bioinformatics: Methods and Protocols (pp. 89–112). Humana Press. 10.1007/978-1-59745-535-0_4

Brzeszcz, J., & Kaszycki, P. (2018). Aerobic bacteria degrading both n-alkanes and aromatic hydrocarbons: an undervalued strategy for metabolic diversity and flexibility. Biodegradation, 29(4), 359–407. 10.1007/s10532-018-9837-x

Buchholz, P. C. F., Feuerriegel, G., Zhang, H., Perez-Garcia, P., Nover, L.-L., Chow, J., Streit, W. R., & Pleiss, J. (2022). Plastics degradation by hydrolytic enzymes: The plastics-active enzymes database-PAZy. Proteins, 90(7), 1443–1456. 10.1002/prot.26325

Deeb, R. A., Hu, H. Y., Hanson, J. R., Scow, K. M., & Alvarez-Cohen, L. (2001). Substrate interactions in BTEX and MTBE mixtures by an MTBE-degrading isolate. Environmental Science & Technology, 35(2), 312–317. 10.1021/es001249j

Duarte, M., Jauregui, R., Vilchez-Vargas, R., Junca, H., & Pieper, D. H. (2014). AromaDeg, a novel database for phylogenomics of aerobic bacterial degradation of aromatics. Database: The Journal of Biological Databases and Curation, 2014, bau118. 10.1093/database/bau118

Ekanayaka, A. H., Tibpromma, S., Dai, D., Xu, R., Suwannarach, N., Stephenson, S. L., Dao, C., & Karunarathna, S. C. (2022). A Review of the Fungi That Degrade Plastic. Journal of Fungi (Basel, Switzerland), 8(8). 10.3390/jof8080772

Gambarini, V., Pantos, O., Kingsbury, J. M., Weaver, L., Handley, K. M., & Lear, G. (2022). PlasticDB: a database of microorganisms and proteins linked to plastic biodegradation. Database: The Journal of Biological Databases and Curation,2022. 10.1093/database/baac008

Gan, Z., & Zhang, H. (2019). PMBD: a Comprehensive Plastics Microbial Biodegradation Database. Database: The Journal of Biological Databases and Curation, 2019. 10.1093/database/baz119

Geyer, R. (2020). Chapter 2 - Production, use, and fate of synthetic polymers. In T. M. Letcher (Ed.), Plastic Waste and Recycling (pp. 13–32). Academic Press. 10.1016/B978-0-12-817880-5.00002-5

Ghosal, D., Ghosh, S., Dutta, T. K., & Ahn, Y. (2016). Current State of Knowledge in Microbial Degradation of Polycyclic Aromatic Hydrocarbons (PAHs): A Review. Frontiers in Microbiology, 7, 1369. 10.3389/fmicb.2016.01369

Gu, Z., Eils, R., & Schlesner, M. (2016). Complex heatmaps reveal patterns and correlations in multidimensional genomic data. Bioinformatics, 32(18). 10.1093/bioinformatics/btw313

Hajighasemi, M., Tchigvintsev, A., Nocek, B., Flick, R., Popovic, A., Hai, T., Khusnutdinova, A. N., Brown, G., Xu, X., Cui, H., Anstett, J., Chernikova, T. N., Brüls, T., Le Paslier, D., Yakimov, M. M., Joachimiak, A., Golyshina, O. V., Savchenko, A., Golyshin, P. N., … Yakunin, A. F. (2018). Screening and Characterization of Novel Polyesterases from Environmental Metagenomes with High Hydrolytic Activity against Synthetic Polyesters. Environmental Science & Technology, 52(21), 12388–12401. 10.1021/acs.est.8b04252

Horton, A. A. (2022). Plastic pollution: When do we know enough? Journal of Hazardous Materials, 422, 126885. 10.1016/j.jhazmat.2021.126885

Hua, F., & Wang, H. Q. (2014). Uptake and trans-membrane transport of petroleum hydrocarbons by microorganisms. Biotechnology, Biotechnological Equipment, 28(2), 165–175. 10.1080/13102818.2014.906136

Huerta-Cepas, J., Szklarczyk, D., Heller, D., Hernández-Plaza, A., Forslund, S. K., Cook, H., Mende, D. R., Letunic, I., Rattei, T., Jensen, L. J., von Mering, C., & Bork, P. (2019). eggNOG 5.0: a hierarchical, functionally and phylogenetically annotated orthology resource based on 5090 organisms and 2502 viruses. Nucleic Acids Research, 47(D1), D309–D314. 10.1093/nar/gky1085

Ji, Y., Mao, G., Wang, Y., & Bartlam, M. (2013). Structural insights into diversity and n-alkane biodegradation mechanisms of alkane hydroxylases. Frontiers in Microbiology, 4, 58. 10.3389/fmicb.2013.00058

Jones, P., Binns, D., Chang, H.-Y., Fraser, M., Li, W., McAnulla, C., McWilliam, H., Maslen, J., Mitchell, A., Nuka, G., Pesseat, S., Quinn, A. F., Sangrador-Vegas, A., Scheremetjew, M., Yong, S.-Y., Lopez, R., & Hunter, S. (2014). InterProScan 5: genome-scale protein function classification. Bioinformatics, 30(9), 1236–1240. 10.1093/bioinformatics/btu031

Kaczorek, E., Pacholak, A., Zdarta, A., & Smulek, W. (2018). The Impact of Biosurfactants on Microbial Cell Properties Leading to Hydrocarbon Bioavailability Increase. Colloids and Interfaces, 2(3), 35. 10.3390/colloids2030035

Kaushal, J., Khatri, M., & Arya, S. K. (2021). Recent insight into enzymatic degradation of plastics prevalent in the environment: A mini - review. Cleaner Engineering and Technology, 2, 100083. 10.1016/j.clet.2021.100083

Kavitha, R., & Bhuvaneswari, V. (2021). Assessment of polyethylene degradation by biosurfactant producing ligninolytic bacterium. Biodegradation, 32(5), 531–549. 10.1007/s10532-021-09949-8

Khot, V., Zorz, J., Gittins, D. A., Chakraborty, A., Bell, E., Bautista, M. A., Paquette, A. J., Hawley, A. K., Novotnik, B., Hubert, C. R. J., Strous, M., & Bhatnagar, S. (2021). CANT-HYD: A Curated Database of Phylogeny-Derived Hidden Markov Models for Annotation of Marker Genes Involved in Hydrocarbon Degradation. Frontiers in Microbiology, 12, 764058. 10.3389/fmicb.2021.764058

Krieger, C. J., Zhang, P., Mueller, L. A., Wang, A., Paley, S., Arnaud, M., Pick, J., Rhee, S. Y., & Karp, P. D. (2004). MetaCyc: a multiorganism database of metabolic pathways and enzymes. Nucleic Acids Research, 32(Database issue), D438–D442. 10.1093/nar/gkh100

Lechner, M., Findeiss, S., Steiner, L., Marz, M., Stadler, P. F., & Prohaska, S. J. (2011). Proteinortho: detection of (co-)orthologs in large-scale analysis. BMC Bioinformatics, 12, 124. 10.1186/1471-2105-12-124

Leinonen, R., Akhtar, R., Birney, E., Bower, L., Cerdeno-Tárraga, A., Cheng, Y., Cleland, I., Faruque, N., Goodgame, N., Gibson, R., Hoad, G., Jang, M., Pakseresht, N., Plaister, S., Radhakrishnan, R., Reddy, K., Sobhany, S., Ten Hoopen, P., Vaughan, R., … Cochrane, G. (2011). The European Nucleotide Archive. Nucleic Acids Research, 39(Database issue), D28–D31. 10.1093/nar/gkq967

Madzak, C. (2021). Yarrowia lipolytica Strains and Their Biotechnological Applications: How Natural Biodiversity and Metabolic Engineering Could Contribute to Cell Factories Improvement. Journal of Fungi (Basel, Switzerland), 7(7). 10.3390/jof7070548

Moreno, R., & Rojo, F. (2017). Enzymes for aerobic degradation of alkanes in bacteria. In Aerobic Utilization of Hydrocarbons, Oils and Lipids (pp. 1–25). Springer International Publishing. 10.1007/978-3-319-39782-5_6-1

Muriel-Millán, L. F., Millán-López, S., & Pardo-López, L. (2021). Biotechnological applications of marine bacteria in bioremediation of environments polluted with hydrocarbons and plastics. Applied Microbiology and Biotechnology, 105(19), 7171–7185. 10.1007/s00253-021-11569-4

Murphy, C. D., Quirke, S., & Balogun, O. (2008). Degradation of fluorobiphenyl by Pseudomonas pseudoalcaligenes KF707. FEMS Microbiology Letters, 286(1), 45–49. 10.1111/j.1574-6968.2008.01243.x

Ogata, H., Goto, S., Sato, K., Fujibuchi, W., Bono, H., & Kanehisa, M. (1999). KEGG: Kyoto Encyclopedia of Genes and Genomes. Nucleic Acids Research, 27(1), 29–34. 10.1093/nar/27.1.29

Oliveira, J. S., Araújo, W., Lopes Sales, A. I., Brito Guerra, A. de, Silva Araújo, S. C. da, de Vasconcelos, A. T. R., Agnez-Lima, L. F., & Freitas, A. T. (2015). BioSurfDB: knowledge and algorithms to support biosurfactants and biodegradation studies. Database: The Journal of Biological Databases and Curation, 2015. 10.1093/database/bav033

Pathak, U., Jhunjhunwala, A., Singh, S., Bajaj, N., & Mandal, T. (2022). Chapter 19 - Potentiality of enzymes as a green tool in degradation of petroleum hydrocarbons. In P. Das, S. Manna, & J. K. Pandey (Eds.), Advances in Oil-Water Separation (pp. 337–351). Elsevier. 10.1016/B978-0-323-89978-9.00024-0

Phale, P. S., Malhotra, H., & Shah, B. A. (2020). Chapter One - Degradation strategies and associated regulatory mechanisms/features for aromatic compound metabolism in bacteria. In G. M. Gadd & S. Sariaslani (Eds.), Advances in Applied Microbiology (Vol. 112, pp. 1–65). Academic Press. 10.1016/bs.aambs.2020.02.002

Prenafeta-Boldú, F. X., de Hoog, G. S., & Summerbell, R. C. (2018). Fungal Communities in Hydrocarbon Degradation. In T. J. McGenity (Ed.), Microbial Communities Utilizing Hydrocarbons and Lipids: Members, Metagenomics and Ecophysiology (pp. 1–36). Springer International Publishing. 10.1007/978-3-319-60063-5_8-1

Prince, R. C., Gramain, A., & McGenity, T. J. (2010). Prokaryotic Hydrocarbon Degraders. In K. N. Timmis (Ed.), Handbook of Hydrocarbon and Lipid Microbiology (pp. 1669–1692). Springer Berlin Heidelberg. 10.1007/978-3-540-77587-4_118

Ru, J., Huo, Y., & Yang, Y. (2020). Microbial Degradation and Valorization of Plastic Wastes. Frontiers in Microbiology, 11, 442. 10.3389/fmicb.2020.00442

Shimao, M. (2001). Biodegradation of plastics. Current Opinion in Biotechnology, 12(3), 242–247. 10.1016/s0958-1669(00)00206-8

Shrivastava, A. (2018). Introduction to Plastics Engineering. Elsevier Science. https://play.google.com/store/books/details?id=yM60tAEACAAJ

Soberón-Chávez, G. (2010). Biosurfactants: From Genes to Applications. Springer Berlin Heidelberg. 10.1007/978-3-642-14490-5

Soberón-Chávez, G., González-Valdez, A., Soto-Aceves, M. P., & Cocotl-Yañez, M. (2021). Rhamnolipids produced by Pseudomonas: from molecular genetics to the market. Microbial Biotechnology, 14(1), 136–146. 10.1111/1751-7915.13700

Srikanth, M., Sandeep, T. S. R. S., Sucharitha, K., & Godi, S. (2022). Biodegradation of plastic polymers by fungi: a brief review. Bioresources and Bioprocessing, 9(1), 1–10. 10.1186/s40643-022-00532-4

Vimala, P. P., & Mathew, L. (2016). Biodegradation of polyethylene using Bacillus subtilis. Procedia Technology, 24, 232–239. 10.1016/j.protcy.2016.05.031

Wentzel, A., Ellingsen, T. E., Kotlar, H.-K., Zotchev, S. B., & Throne-Holst, M. (2007). Bacterial metabolism of long-chain n-alkanes. Applied Microbiology and Biotechnology, 76(6), 1209–1221. 10.1007/s00253-007-1119-1

Widdel, F., & Musat, F. (2010). Diversity and Common Principles in Enzymatic Activation of Hydrocarbons. In K. N. Timmis (Ed.), Handbook of Hydrocarbon and Lipid Microbiology (pp. 981–1009). Springer Berlin Heidelberg. 10.1007/978-3-540-77587-4_70

Xu, X., Liu, W., Tian, S., Wang, W., Qi, Q., Jiang, P., Gao, X., Li, F., Li, H., & Yu, H. (2018). Petroleum Hydrocarbon-Degrading Bacteria for the Remediation of Oil Pollution Under Aerobic Conditions: A Perspective Analysis. Frontiers in Microbiology, 9, 2885. 10.3389/fmicb.2018.02885

Yoshida, T., Sada, Y., & Nagasawa, T. (2010). Bioconversion of 2,6-dimethylpyridine to 6-methylpicolinic acid by Exophiala dermatitidis (Kano) de Hoog DA5501 cells grown on n-dodecane. Applied Microbiology and Biotechnology, 86(4), 1165–1170. 10.1007/s00253-009-2419-4

Zhang, J., Chen, J., Jia, R., Dun, Z., Wang, B., Hu, X., & Wang, Y. (2018). Selection and evaluation of microorganisms for biodegradation of agricultural plastic film. 3 Biotech, 8(7), 308. 10.1007/s13205-018-1329-5

